# MorphoHub: A Platform for Petabyte-Scale Multi-Morphometry Generation

**DOI:** 10.1101/2021.01.09.426010

**Authors:** Shengdian Jiang, Yimin Wang, Lijuan Liu, Sujun Zhao, Mengya Chen, Xuan Zhao, Peng Xie, Liya Ding, Zongcai Ruan, Hong-Wei Dong, Giorgio A. Ascoli, Michael Hawrylycz, Hongkui Zeng, Hanchuan Peng

**Affiliations:** SEU-ALLEN Joint Center, Institute for Brain and Intelligence, Southeast University, Nanjing, Jiangsu, China; School of Computer Science and Engineering, Southeast University, Nanjing, Jiangsu, China; School of Computer Engineering and Science, Shanghai University, Shanghai, China; Department of Neurobiology, David Geffen School of Medicine at UCLA, Los Angeles, CA 90095, USA; Volgenau School of Engineering, George Mason University, Fairfax, VA 22030, USA; Allen Institute for Brain Science, Seattle, WA 98109, USA

## Abstract

Recent advances in neuroscience make the extraction of full neuronal morphology at whole brain dataset available. To produce quality morphometry at large scale, it is highly desirable but extremely challenging to efficiently handle petabyte-scale high-resolution whole brain imaging database. Here, we developed a multi-level method to produce high quality somatic, dendritic, axonal, and potential synaptic morphometry, which was made possible by utilizing necessary petabyte hardware and software platform to optimize both the data and workflow management. Our method also boosts data sharing and remote collaborative validation. We highlight a petabyte application dataset involving 62 whole mouse brains, from which we identified 50,233 somata of individual neurons, profiled the dendrites of 11,322 neurons, reconstructed the full 3-D morphology of more than one thousand neurons including their dendrites and full axons, and detected million scale putative synaptic sites derived from axonal boutons. Analysis and simulation of these data indicate the promise of this approach for modern large-scale morphology applications.

## RESULTS

We attempted to meet the petabyte (PB)-scale whole-brain computing challenge by introducing a technology that involves a hardware platform that is able to handle petabytes of data storage, sharing, computing and visualization, a software platform called *MorphoHub* that can utilize such hardware platform, and most importantly, a mechanism to scale up the synergized automated computation and multi-user collaboration for effective validation and correction. Our method is centered around reconstructing the multi-level information of single neuron’s morphology (**Fig. 1a**) at the whole-brain scale for a number of brains. The *MorphoHub* software package is able to streamline the workflow of imaging data management, visualization, reconstruction and collaboration, and data sharing (**Fig. 1b**, **Methods**). Using this approach, we extended several state-of-the-art methods to the PB-scale (**Supplementary Table 1**) and produced multi-morphometry data from such massive image database (**Fig. 1c**). Our method allows a smooth transition from manual and interactive morphometry acquisition to increasingly routine work done by automatic algorithms as we show below.

**Fig 1.**
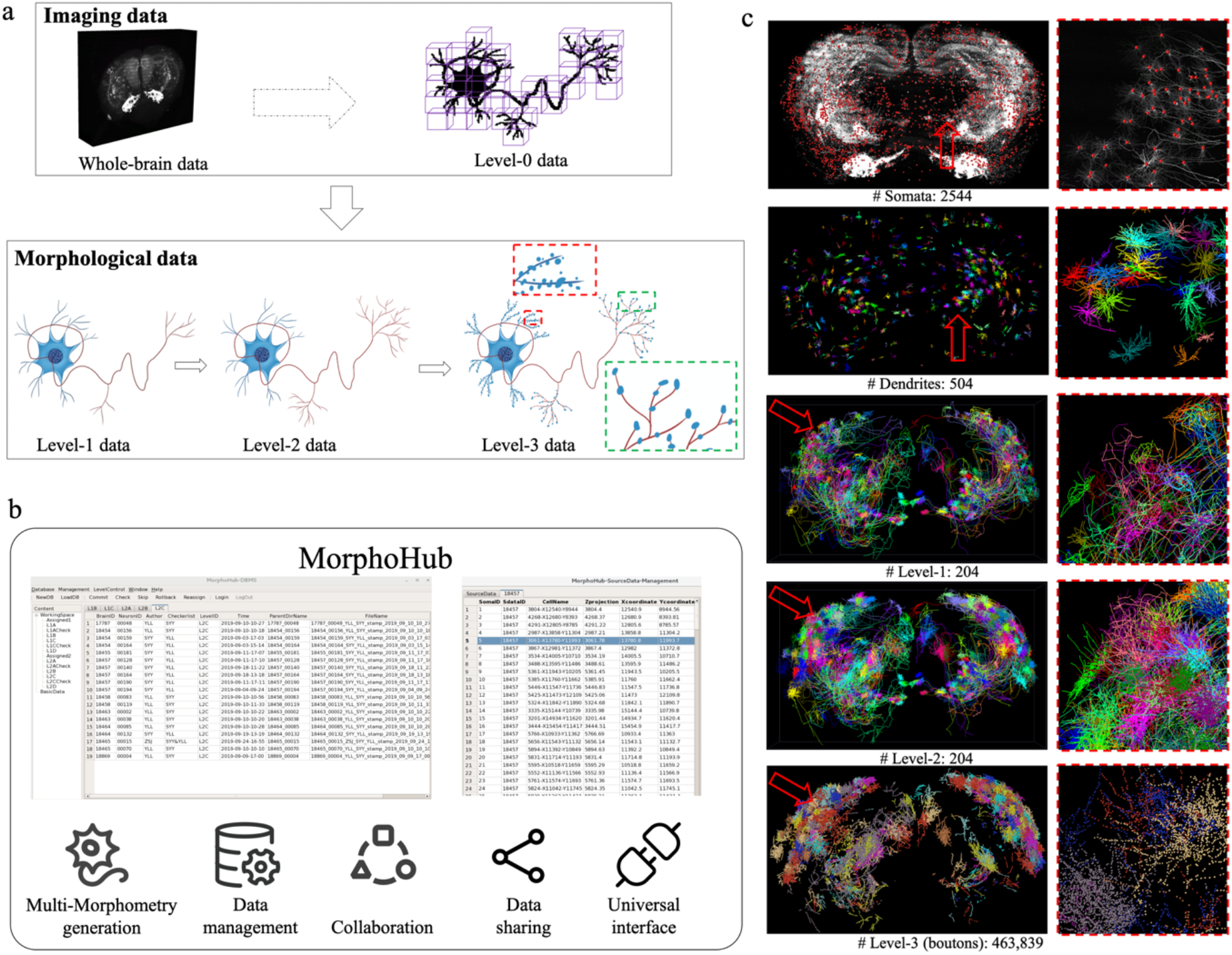
Multi-morphometry data generation from whole-brain imaging datasets. **a** An illustration of the multi-level reconstruction approach. From a whole-brain image containing trillions of voxels (top left), the Level-1 (L1), Level-2 (L2), and Level-3 (L3) data are reconstructed in sequence (bottom). Moreover, a concise Level-0 (L0) imaging data is also generated based on the reconstructed morphometry (top right). **b** The MorphoHub system for the generation of multi-morphometry data, management and visualization of all related data and workflow, data sharing and extended functions. **c** Examples of the multi-morphometry data reconstructed from one Brain (Brain id: 18454). From top to bottom are the somata, dendrites, L1, L2, and L3 data, respectively, with zoom-in panels for red arrows shown on the right.

A key component of our method is a three-level (L1, L2, and L3) reconstruction approach (**Fig. 1a**). We incrementally reconstruct morphological components of neurons, including somata, dendrites, axons, spines and boutons, *only* when such information can be produced faithfully and affordably. Specifically, an L1 reconstruction contains the full dendritic arbor and the skeleton of all axonal neurite tracts, excluding the fine structures of distal axonal arbors (**Fig. 1a**). An L2 reconstruction contains the complete structures of all neuronal arbors (**Fig. 1a**). An L3 reconstruction contains the identification of two key elements of synaptic connectivity, dendritic spines and axonal boutons, as well as other structures of potential interest (e.g., specific topology of axonal branching patterns, modeling of specific neuronal compartments’ shape) (**Fig. 1a**).

The proposed multi-level reconstruction method is generic and scalable to single neuron datasets of arbitrary size if proper data structure and data workflow are in place. To provide such capability for a real PB-scale computing environment, we developed *MorphoHub* to manage all data flow and processing procedures in an integrated way (**Fig. 1b** and **Supplementary Fig. 1**, **Methods**). *MorphoHub* handles four heterogeneous data types, including image volumes, neuron morphology, meta-data of user interactions, and data management (conversion, storage, transferring/sharing) schemas, for a PB-scale database. We also engineered a universal application interface (**Fig. 1b** and **Supplementary Fig. 2**) in *MorphoHub* so that it could invoke additional image analysis and validation tools when needed.

To demonstrate the capability of this approach, we built an image database called *D62* consisting of 62 whole mouse brain images. *D62* has in total 713.35 teravoxels, 1.43 petabytes in native image space, and 973 terabytes in compressed file space (**Supplementary Data 1**). *MorphoHub* ran smoothly on *D62* and allowed us to precisely pinpoint somata of 50,233 neurons using TeraFly (100% accuracy validated by independent annotators). For each neuron, we then reconstructed the dendrites automatically, followed by feature-based screening and visual validation (**Fig. 1c**, **Methods**). Using this workflow, we produced traceable dendritic results of 11,322 neurons. Due to the scale of the problem, similar results were hard to obtain using other software.

All multi-morphometry data produced were also registered to the Allen Common Coordinate Framework (CCF)^37^ to see how the distributions of each data level correlate with others (**Fig. 2**). In this way, various brain regions (the white colored regions), dendrites, axons, and boutons can all be identified (**Fig. 2a-e**). As a result, we output a summary matrix with rows representing neurons, columns representing a unique CCF parcel, and numerical entries expressing, for each neuron and corresponding parcel, the axonal and dendritic length, the number of boutons, and the (binary) presence of soma. Such representation lends itself to highly informative quantitative analyses, such as pairwise probability of directional connection between neurons (dot product of presynaptic axonal values and postsynaptic dendritic values) and projection similarity (arccosine distance between axonal values of two neurons). Fig. 2 shows that the distributions of these neuronal entities do not correlate globally. Instead, they exhibit regional enrichment of which the pattern is hard to observe when only local brain areas are analyzed.

**Fig 2.**
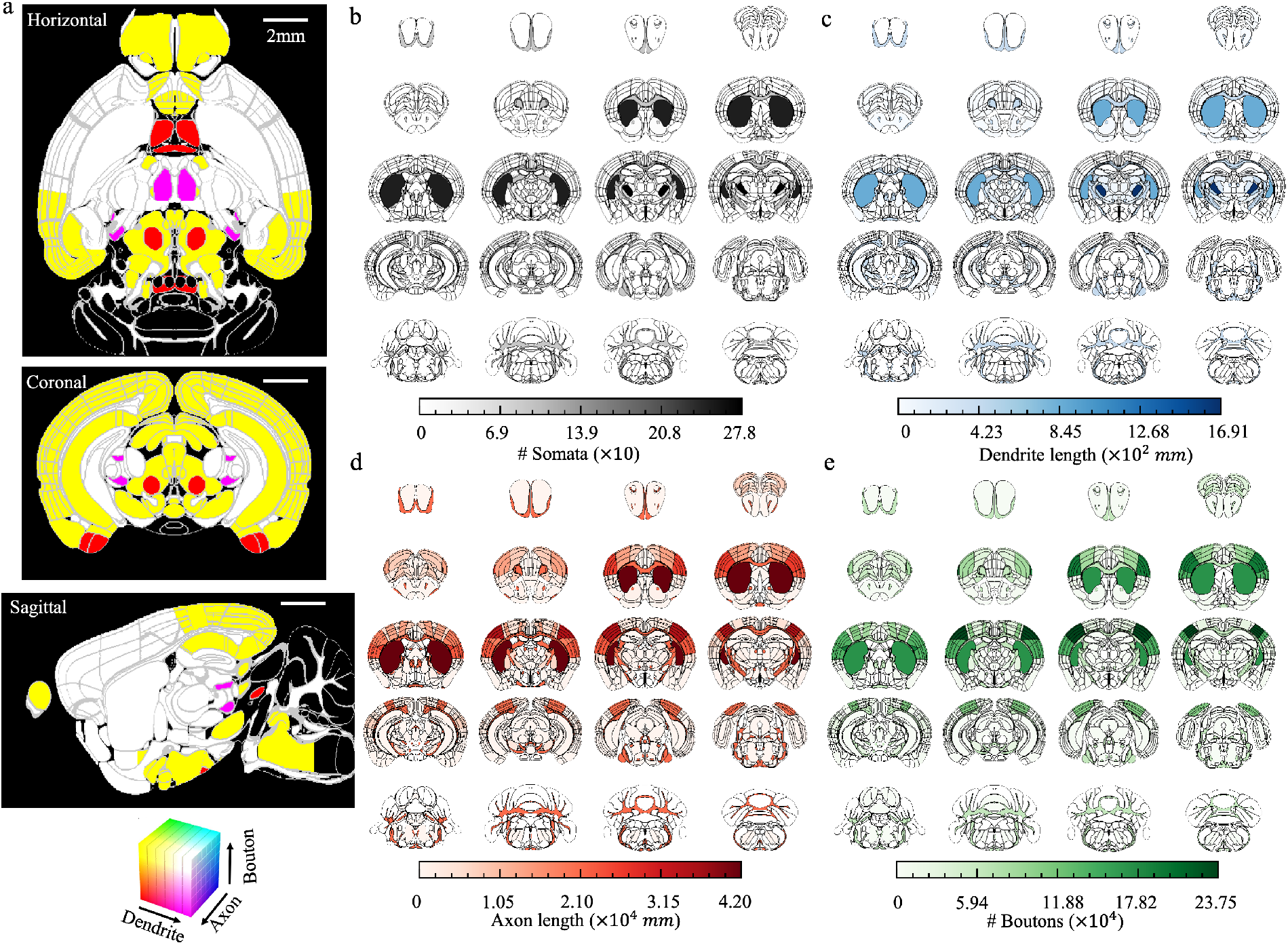
Visualization of the multi-morphometry data in R1050. **a** The color-coded joint distribution of dendrites, axons, and boutons. From top to bottom: horizontal view (slice no. 165), coronal view (slice no. 335), and sagittal view (slice no. 148). Colors indicate the densities of dendrites, axons, and boutons normalized to the standard RGB color space. Scale bar: 2mm. **b** Individual distributions of somata in R1050. Each inset corresponds to the combination of 25 consecutive coronal slices, in which the brain regions were colored according to the densities of somata. The darker the color, the higher the density. **c-e** Similar visualizations for dendrites, axons, and boutons in R1050, respectively.

For PB-scale computing, the speed of data I/O for storage and data sharing across networks (Internet or intranets) becomes more critical than in applications at small scale. It is essential to reduce data volume without compromising visualization and analysis of such large data. We observed that an L1-L2-L3 morphology trio of a neuron will always be sparse and that the spine and bouton information in the L3 data could be described using a neighborhood around the neuron skeleton in preceding levels. To utilize this observation, we developed a compact L0 representation of a neuron for effective imaging data management, sharing, and computation (**Fig. 3a**, **Methods**). The key idea is that the L0 data of a neuron represents a tightly bounded image region that covers the L1-L2-L3 trio area. Because dendritic spines typically attach dendritic fiber orthogonally, and axonal boutons scatter along axons, for any given L2 (or L1) skeleton we conveniently extracted a series of piecewise 512×512×512-voxel 3-D image-tiles around the skeleton to cover all spines and boutons and at the same time to allow fast image file I/O. In this way, an L0 representation identifies an image sub-region that contains all parts of a specific neuron.

**Fig 3.**
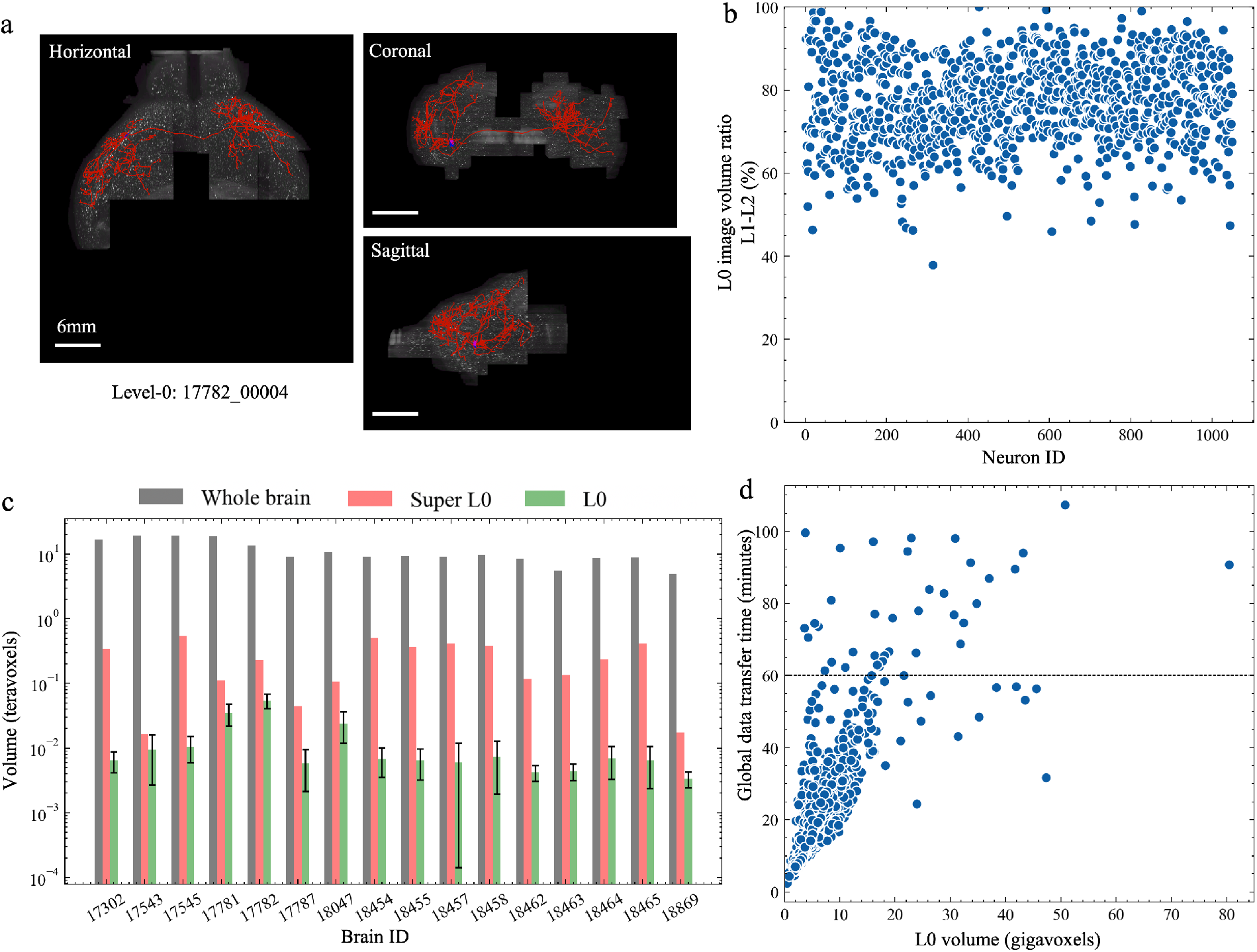
The L0 representation of imaging data. **a** The L0 image (shown in horizontal, coronal, and sagittal views) for neuron 17782_00004, overlaid with its L2 reconstruction. **b** For neurons in R1050, the ratio of L0 image volume generated from L1 data over that generated from L2 data. **c** Comparisons of the size of whole-brain images, the average size of L0 data, and the size of the “super L0-data” (union of all L0 data of neurons). Error bar: SD. **d** Time for transferring 1050 L0 images between two research centers in Asia (SEU-ALLEN) and America (BIL).

For *R1050* reconstructed from *D62*, on average the L0 data generated based on an L1 skeleton contains more than 77% of that generated based on the corresponding L2 reconstruction (**Fig. 3b**). In addition, the L0 data of a neuron typically occupies three orders of magnitude less image volume compared to the whole brain image (**Fig. 3c**). The L0 data of the largest neuron in this work has ~80 gigavoxels, while the mean value and standard deviation of the volume of L0 data of 1,050 fully reconstructed neurons are 6.75 and 5.94 gigavoxels, respectively (**Fig. 3d, Supplementary Table 2**). Practically, even the union of all L0 data of neurons, denoted as Super L0, in a sparsely labeled brain still has 1 to 3 orders of magnitude fewer voxels compared to the total volume of the brain (**Fig. 3c**). In this way, the multi-morphometry framework allows thousands of fold better efficiency in both storing and transferring the essential image data and quantitative shape information of neurons. This utility greatly simplifies the previously challenging data sharing task. Indeed, without accelerated content delivery, currently it is possible to transfer the L0 data in *R1050* between the data production center (SEU-ALLEN) in Asia and one data releasing facility (BICCN Image Library, Pittsburgh supercomputing center) in North America (**Fig. 3d**). This direct data sharing replaced a previous way to bulk ship hard drives containing the massive amount data back-and-forth across continents.

With the L0-L1-L2-L3 quadruple data, we further enhanced the scale and faithfulness of our multi-morphometry produced in two ways. First, since the L0 data has a much smaller volume and thus is much easier to share across network, we developed a collaborative mode that interconnects a number of formerly autonomous TeraFly/TeraVR users to synergistically work on the L0 data directly (**Supplementary Fig. 3**). This method parallelizes the workflow and thus improves the data production rate. The cooperative work of multiple annotators also elevates the faithfulness of the resultant morphometry. Second, we used a deep learning network to learn from the L0-L*x* (*x* = 1,2,3) pair. The trained model was used to detect specific neuronal features, such as neurite skeleton or axonal terminals (**Supplementary Fig. 4**). This process can be repeatedly optimized based on progressively more and more accurate L0-L1-L2-L3 quadruple data. Such automation also increases the data production rate for PB-scale computing.

## DISCUSSION

In this study, we demonstrated a robust PB-scale informatics platform to generate large-scale single neuron reconstructions suitable for multi-scale biological analysis. Our approach has several advantages: (1) efficient multi-level production and management of whole-brain neuron reconstructions; (2) conducting morphological analysis and cell typing globally and at multi-resolution; (3) enabling the investigation of the convergence or divergence of neuronal projections by analyzing distribution of neuronal arbors across brain regions; (4) comparison of various neuronal elements and sub-structures with respect to the types of cells. Taken altogether, our whole-brain multi-morphometry approach provides a framework to produce hierarchical datasets that synchronize brain anatomy, single neuron morphology, sub-neuronal structures, and potential pre-synaptic sites, all mapped onto a standardized atlas. Our method will be useful for further studies of neuronal circuits based on whole-brain imaging, not only for mouse brains but also for other model systems such as monkey brains.

Our work furthers previous effort to use light microscopy (LM) to visualize and detect synapses around neurite tracts labeled by genetic markers^1,2^ or antibodies^3^, which were limited to partial neuronal structures in local brain regions. In this study we used fMOST data as a showcase and for putative synaptic sites we have focused on axonal boutons. As a generic computational framework, our approach is applicable to various datasets produced with different methods and collected with different imaging modalities.

The multi-level reconstruction approach is being enhanced in various ways. In addition to various workstation/PC clients, virtual reality consoles, super-computing, and big-display walls that are already integrated in our software *MorphoHub*, mobile applications (APPs) for more intelligent and automated neuron tracing are being developed and added onto our software. We are also deploying *MorphoHub* for data servers in the cloud and scaling up the capability for concurrent data serving of distributed users. We hope these engineering efforts would lead to a new globally accessible platform that has potential to bring the current productivity to the next level, especially addressing challenges in completing neuron morphometry more efficiently, producing more fine-scale morphometry such as synapses with their shapes and statistics, integrating more automation through the use of AI, sharing of imaging data remotely at affordable cost, and international collaboration of neuroanatomists and other interested users.

## CONCLUSION

Neuronal morphology is an essential component of cell type identity in the brain and an essential determinant of connectivity and circuit function. Large scale accurate neuronal profiling necessitates advanced methods in computational processing to effectively manage storage and bandwidth for collaborative segmentation and annotation. The data in this study shows our petabyte-scale computing framework is able to provide a solution to modern anatomic workflow requirements that are now demanded for very large-scale morphometry.

## DATA AND CODE AVAILABILITY

The whole brain image datasets are released under BICCN’s Brain Image Library (BIL) at Pittsburgh Supercomputing Center.

## ACKNOWLEDGMENTS

This work is funded by Southeast University (SEU) to support Open Science collaboration. SEU supported the development of the methods and informatics data management and analysis pipeline at the SEU-Allen Joint Center. We are grateful to the Allen Institute for imaging datasets collected through multiple grant awards from institutes under the National Institutes of Health (NIH), including award number R01EY023173 from The National Eye Institute to H.Z., U01MH105982 from the National Institute of Mental Health and Eunice Kennedy Shriver National Institute of Child Health & Human Development to H.Z., and U19MH114830 from the National Institute Of Mental Health to H.Z. The content is solely the responsibility of the authors and does not necessarily represent the official views of NIH and its subsidiary institutes. The Allen Institute affiliated authors wish to thank the Allen Institute founder, Paul G. Allen, for his vision, encouragement, and support. G.A.A. acknowledges NIH grants R01NS39600, U01MH14829, and R01NS86082. We thank Zhi Zhou, Yuanyuan Song, Lulu Yin, Shichen Zhang, Jintao Pan, Yanting Liu, Guodong Hong, Jia Yuan, Yanjun Duan, Yaping Wang, Qiang Ouyang, Zijun Zhao, Wan Wan, Peng Wang, Ping He, Lingsheng Kong, Feng Xiong, and other team mmebers for the support of data production.

## AUTHOR CONTRIBUTIONS

H.P. conceptualized the study, envisioned and led the development of this platform and various analyses. S.J., and Y.W. co-developed the *MorphoHub* system and the multi-level neuron reconstruction protocols. Y.W. provided the support of *TeraVR*. M.C. led the identification of somata in D62. S.Z., X.Z. assisted in *MorphoHub* development. P.X. and L.D. assisted in data analysis. Z.R. and H.P. constructed the hardware platform for this study. H.Z. provided the raw imaging dataset for producing D62 and advised on neurobiology. M.H. advised on morphometry and data management. H.D. and G.A. advised on whole brain neuronal structure analysis.

## METHODS

### Hardware and software platform

Our system is designed for the task of multi-morphometry data generation from petabyte-scale whole-brain imaging database. Built upon customized hardware infrastructure and software tools, the system is capable of organizing whole-brain imaging datasets, generating multi-morphometry data, managing data and workflow, and visualization and analysis of the generated data (**Supplementary Figure 1**).

The hardware infrastructure includes the following parts. VR-equipped annotation workstations are used for data visualization, interactive neuron reconstruction, proofreading, etc. A petabyte-scale storage is configured to store the whole-brain imaging datasets, while the multi-morphometry data is managed using cloud-based storage. A computing cluster is deployed for parallel execution of batch work assignments. Moreover, a wall-mounted display array is also available for monitoring the data generation status. The storage server and the computing cluster are connected with a 100 Gbps wired local area network for peta-scale data storage and parallel computing. The annotation workstations, cloud storage, and monitor system reside in a 10 Gbps local area network.

There are four major software components in the *MorphoHub* software package. The *MorphoHub*-DBMS is the core of the entire system which is responsible for the coordination of the overall data generation process. The DendriteGenerator is in charge for parallel generation of dendritic arbors. The L0Generator is useful for creating compact image representation of the reconstructions. The BoutonGenerator is capable of automatically detection of the synaptic boutons located on axons. These components are developed as plugins of Vaa3D^44,45^, thus making *MorphoHub* cross-platform and deployable on various operating systems.

### *MorphoHub*-DBMS

The *MorphoHub*-DBMS manages the generation of multi-morphometry data by either collaborating teams or automatic routines. All the morphometry data files are organized in the *MorphoHub*-Database, where they follow consistent naming rule and are assigned with unique IDs. The validity of the database is monitored by an error checking routine running in the background. Should there be any issue regarding to the morphometry data files, and error message would be generated and displayed on the screen wall system.

During the multi-level generation of the neuron morphometry data, multiple brains, neurons, annotators are involved. Also, for each neuron, the L1, L2, and L3 data, together with a number of intermediate data levels are produced. The *MorphoHub*-DBMS maintains the correct working state for all the neurons and supports a number of operations, including commit (submitting a reconstruction to a higher level), rollback (sending a reconstruction to a previous level), proofreading (requesting for quality check of the reconstruction), etc. Besides, the *MorphoHub*-DBMS also provides supports for user management and task assignment. All the data are periodically synchronized with private or public cloud-based storages for version control, data sharing and collaboration.

A screen wall display system is connected with *MorphoHub* to present useful information for the ongoing morphometry generation. On the display array, some of the latest reconstructed neurons are displayed; the status of the data repository is tracked; preliminary analysis results are also generated dynamically to provide potential insights of the data.

### Generation of somata and dendritic arbors (DendriteGenerator)

The annotators browsed each whole brain image in *D62* using TeraFly and pinpointed soma locations. 2-D maximum intensity projection (MIP) images were generated for each soma-centered region for further validation. Then, centering at each identified soma, a local image volume of 1024×1024×512 was cropped from the from highest resolution of whole brain image. Next, the APP2^46^ algorithm was invoked for the tracing of dendritic arbors. In particular, a number of background thresholds (15, 20, ..., 40) were adopted for each tracing routine. The tasks were submitted to the cluster server for parallel computation. The results were retrieved and stored only if the execution time was under 30 seconds. Then, we leveraged the gold-standard datasets, e.g., a set of manually annotated and validated dendritic arbors, to form rules for further screening the automatic reconstruction results. The [min, max] interval of the following five features of the dendritic arbors were considered, including ‘Tips’, ‘Length’, ‘Max Path Distance’, ‘Average Bifurcation Angle Remote’, and ‘Max Branch Order’. An automatic tracing result qualified if more than four features conformed with the gold-standard. In case more than one tracing qualified for a certain soma location, the result with larger overall tracing length was selected. In visual screening, tracing results were removed when multiple cells were connected.

### Generation of the L0 imaging data (L0Generator)

The L0 imaging data is generated based on the corresponding morphometry data, e.g., L1 data, L2 data, or even a dendritic arbor. The L0 data contains the image regions that cover all the anatomical structures of the morphometry and is organized in a TeraFly-compatible hierarchical form, just as the whole-brain imaging data. The approach to generate L0 data is described below. For a given neuron, each fundamental morphological element, i.e., nodes and edges, is examined. The local image block in the whole-brain dataset that contains the element is found and marked as “relevant”. Then, all the “relevant” images are combined into the union set from which the compact L0 data are finally generated by building a multi-resolution image hierarchy.

## SUPPLEMENTARY TABLES

**Supplementary Table 1.**
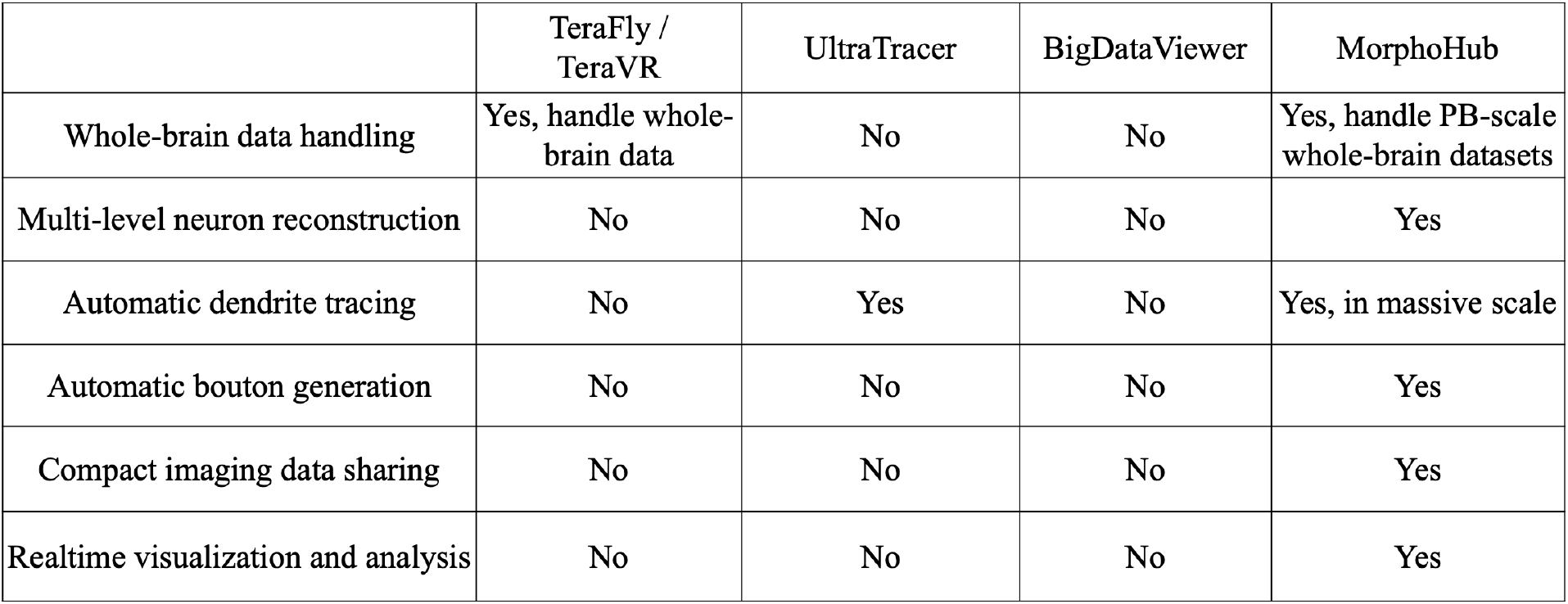
Comparisons between MorphoHub and relevant software for whole mouse-brain imaging data

|  | TeraFly /<br>TeraVR | UltraTracer | BigDataViewer | MorphoHub |
| --- | --- | --- | --- | --- |
| Whole-brain data handling | Yes, handle whole-brain data | No | No | Yes, handle PB-scale whole-brain datasets |
| Multi-level neuron reconstruction | No | No | No | Yes |
| Automatic dendrite tracing | No | Yes | No | Yes, in massive scale |
| Automatic bouton generation | No | No | No | Yes |
| Compact imaging data sharing | No | No | No | Yes |
| Realtime visualization and analysis | No | No | No | Yes |

## SUPPLEMENTARY FIGURES

**Supplementary Figure 1.**
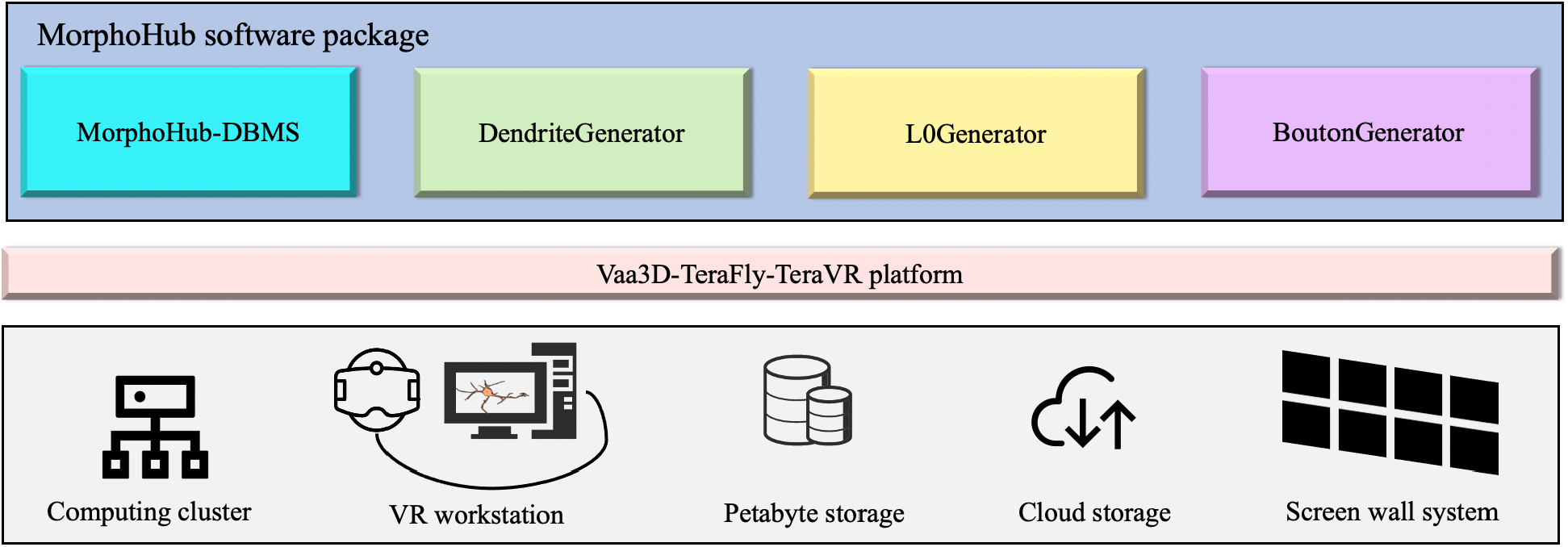
An illustration of our system. A petabyte hardware platform is constructed (bottom row), upon which the MorphoHub software package is developed (upper row). There are four major modules in MorphoHub, which are the MorphoHub-DBMS, DendriteGenerator, L0Generator, and BoutonGenerator, respectively.

**Supplementary Figure 2.**
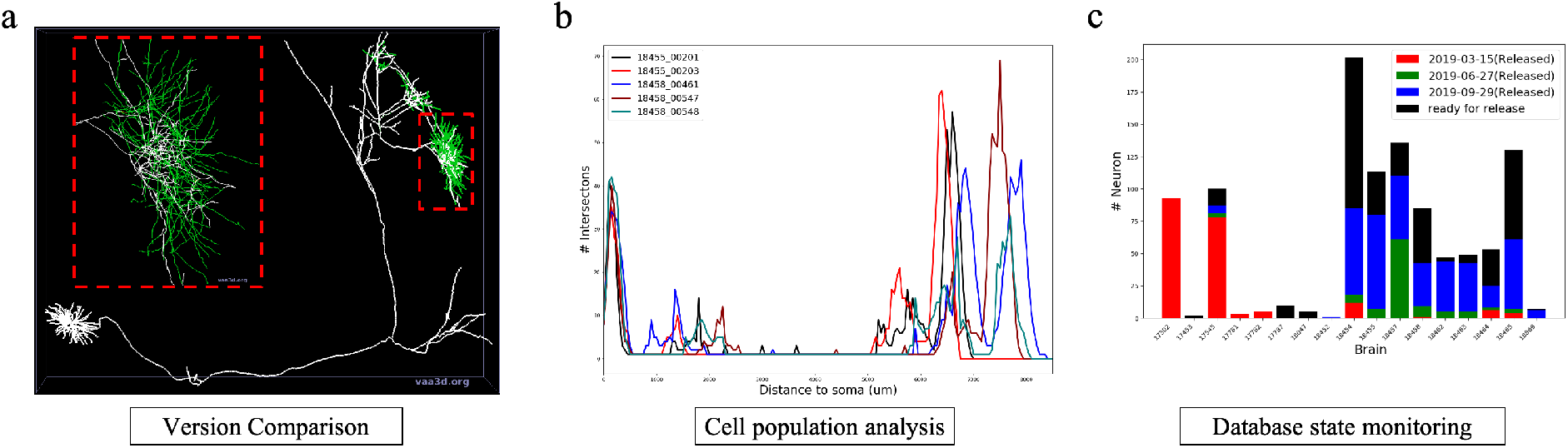
Extended functions via the universal application interface of MorphoHub. **a** Version control tool for comparing the structural difference of two reconstruction versions. (white: identical structures, green: different structures). **b** Sholl analysis for neurons of the same cell type. **c** Monitoring of the morphometry data production. The numbers of generated neurons are shown for each brain and each release date.

**Supplementary Figure 3.**
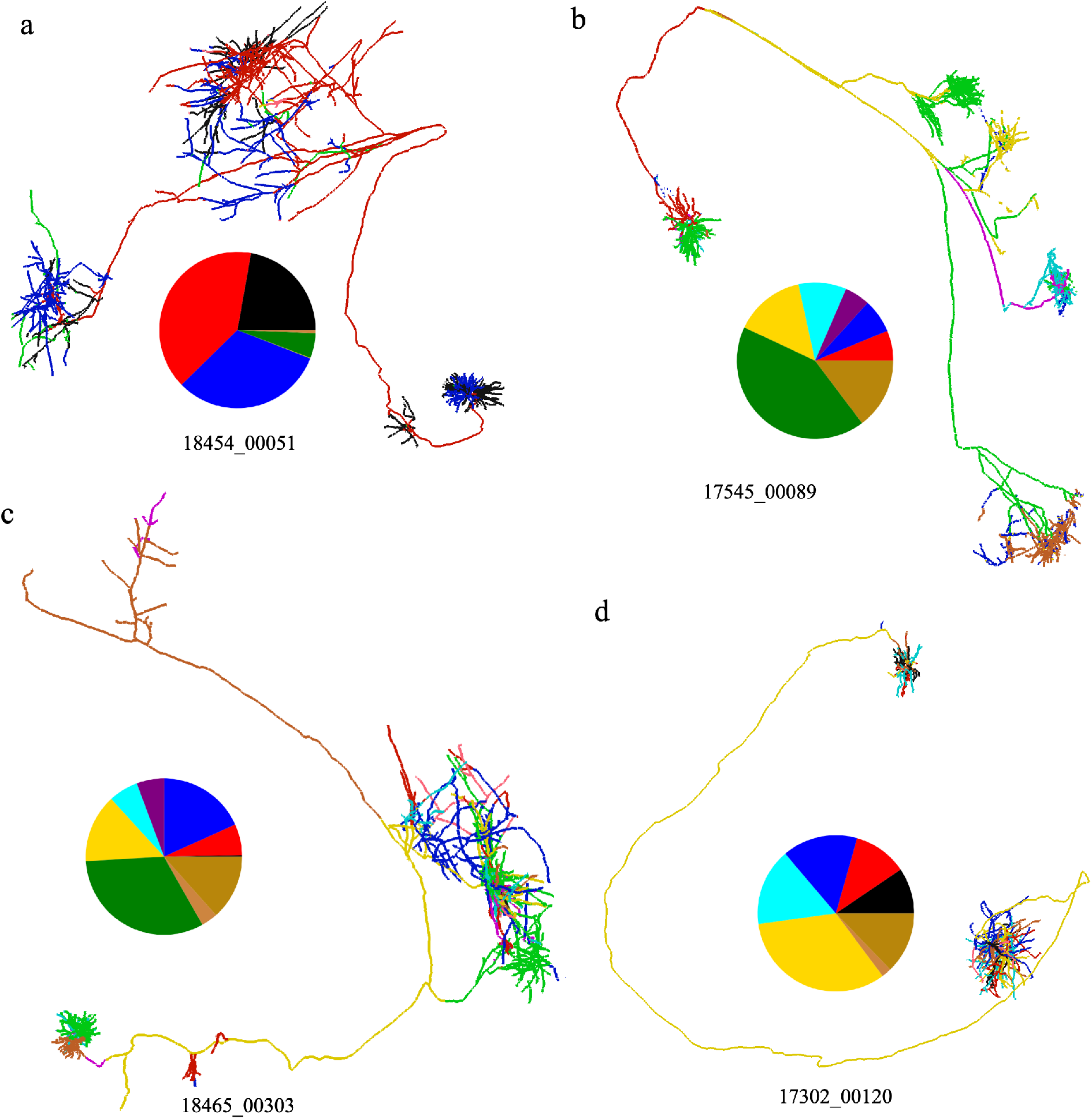
Collaborative neuron reconstruction among several TeraFly/TeraVR clients. **a-d**. L2 neurons that are reconstructed and proofread by a group of annotators. The portion reconstructed by each annotator is represented by a different color. The pie chart corresponds to the reconstruction length of distinct annotators.

**Supplementary Figure 4.**
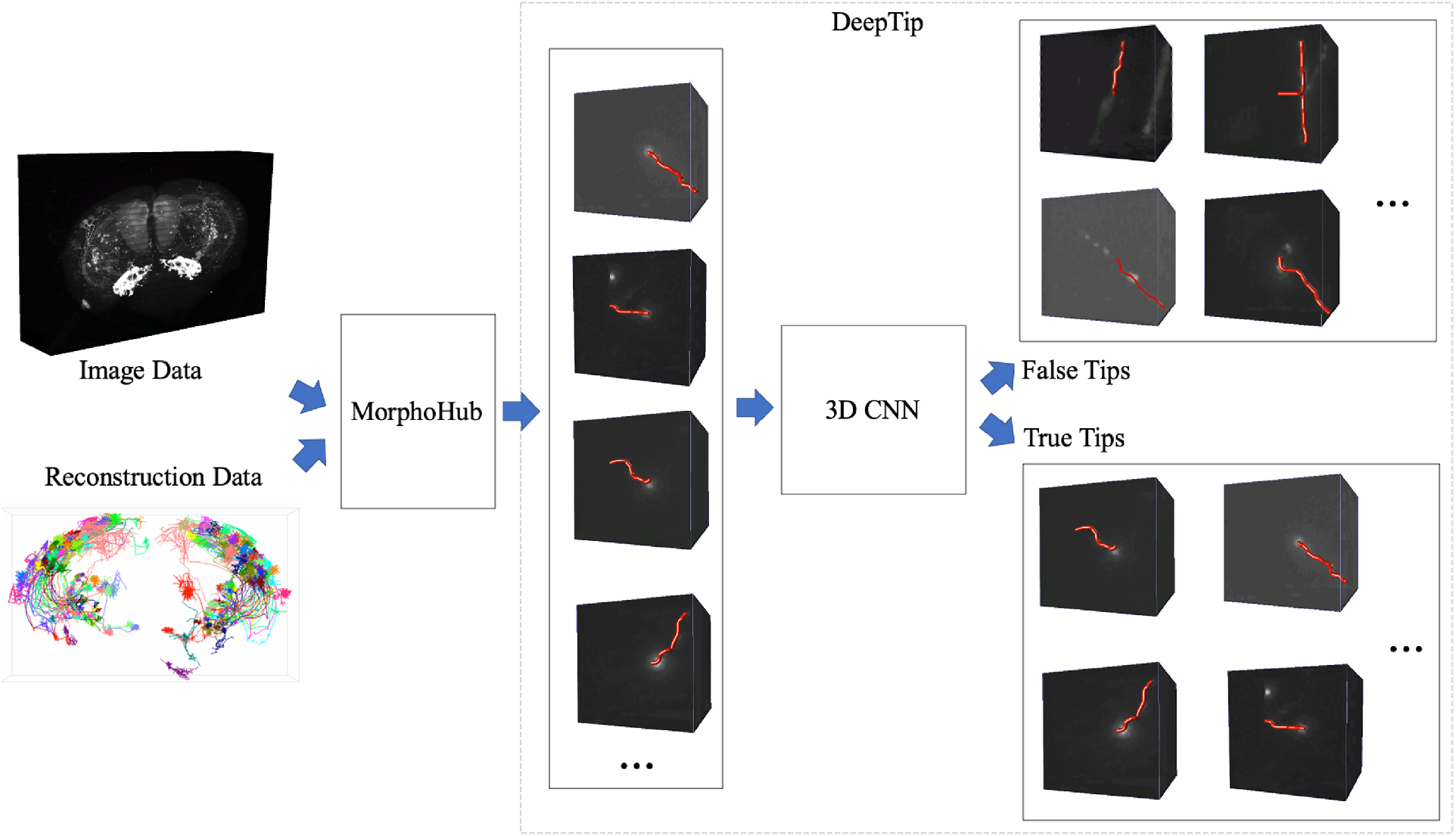
3D-CNN based tip detection. MorphoHub is compatible for integrating various AI components to assist with data production, proofreading and analysis. For example, DeepTip is a deep learning-based component that is integrated in MorphoHub for differentiating true terminal tips of neurites and even automatically correcting the false ones.

