## Supplementary material for "MorphoHub: A Platform for Petabyte-Scale Multi-Morphometry Generation": Fig 1

| Dataset order | Brain ID/Total | Image Size X | Image Size Y | Image Size Z |
|---|---|---|---|---|
| - | (Total) | - | - | - |
| 1 | 17302 | 34,412 | 54,600 | 9,847 |
| 2 | 17543 | 34,748 | 54,600 | 11,333 |
| 3 | 17545 | 35,989 | 54,600 | 10,750 |
| 4 | 17781 | 34,722 | 54,600 | 11,021 |
| 5 | 17782 | 45,000 | 28,000 | 11,786 |
| 6 | 17787 | 36,400 | 24,275 | 11,478 |
| 7 | 18047 | 36,000 | 28,000 | 11,561 |
| 8 | 18452 | 36,400 | 23,423 | 11,028 |
| 9 | 18454 | 35,000 | 26,298 | 11,041 |
| 10 | 18455 | 40,600 | 23,376 | 10,892 |
| 11 | 18457 | 37,800 | 24,275 | 10,955 |
| 12 | 18458 | 40,600 | 25,376 | 10,333 |
| 13 | 18462 | 32,200 | 27,748 | 10,578 |
| 14 | 18463 | 32,200 | 17,478 | 10,842 |
| 15 | 18464 | 35,000 | 26,352 | 10,431 |
| 16 | 18465 | 36,000 | 26,000 | 10,301 |
| 17 | 18869 | 29,400 | 16,915 | 10,923 |
| 18 | 18453 | 39200 | 24400 | 10538 |
| 19 | 17788 | 35000 | 27188 | 11120 |
| 20 | 18459 | 36400 | 22333 | 11264 |
| 21 | 18461 | 28000 | 17838 | 11117 |
| 22 | 18466 | 42000 | 24400 | 11235 |
| 23 | 18467 | 35000 | 25324 | 11389 |
| 24 | 18468 | 37230 | 54600 | 11256 |
| 25 | 18469 | 34412 | 54600 | 10359 |
| 26 | 18470 | 31954 | 54600 | 10226 |
| 27 | 18471 | 39200 | 24400 | 11004 |
| 28 | 18472 | 42000 | 25324 | 10714 |
| 29 | 18671 | 28290 | 37541 | 5018 |
| 30 | 18860 | 42000 | 25324 | 10952 |
| 31 | 18861 | 42000 | 23376 | 10906 |
| 32 | 18862 | 28000 | 17910 | 10798 |
| 33 | 18864 | 27299 | 35001 | 10392 |
| 34 | 18865 | 36401 | 24001 | 11625 |
| 35 | 18866 | 29400 | 19900 | 11374 |
| 36 | 18867 | 40600 | 25376 | 11203 |
| 37 | 18868 | 36401 | 24001 | 10995 |
| 38 | 18053 | 37000 | 25600 | 12500 |
| 39 | 18872 | 26600 | 18449 | 10370 |
| 40 | 236174 | 23814 | 36400 | 12061 |
| 41 | 182722 | 25351 | 37801 | 10725 |
| 42 | 191804 | 18875 | 29401 | 11033 |

| | | | | |
|---|---|---|---|---|
| 43 | 191803 | 17848 | 29401 | 11712 |
| 44 | 191798 | 18893 | 30801 | 10711 |
| 45 | 191797 | 17933 | 30801 | 11197 |
| 46 | 182712 | 25326 | 39201 | 8595 |
| 47 | 191801 | 17865 | 28001 | 11151 |
| 48 | 191807 | 20821 | 30801 | 10830 |
| 49 | 17052 | 34999 | 22900 | 11350 |
| 50 | 17109 | 22793 | 35000 | 10553 |
| 51 | 17300 | 37230 | 54600 | 9954 |
| 52 | 17301 | 33600 | 27188 | 9848 |
| 53 | 17304 | 42194 | 54600 | 10643 |
| 54 | 17539 | 33601 | 23334 | 11143 |
| 55 | 17541 | 39328 | 54600 | 11073 |
| 56 | 17542 | 34748 | 54600 | 11333 |
| 57 | 18049 | 35001 | 30001 | 11412 |
| 58 | 17544 | 39200 | 26351 | 11033 |
| 59 | 17783 | 33600 | 23712 | 10817 |
| 60 | 17785 | 35000 | 25245 | 10509 |
| 61 | 17786 | 36870 | 54600 | 11414 |
| 62 | 18052 | 35000 | 28303 | 10923 |

| #Voxels (TVoxels) | y size (Uncompressec | Size (compressed) (TB) |
|---|---|---|
| 713.35 | 1426.71 | 973.17 |
| 18.50 | 37.00 | 25.24 |
| 21.50 | 43.00 | 29.33 |
| 21.12 | 42.25 | 28.82 |
| 20.89 | 41.79 | 28.50 |
| 14.85 | 29.70 | 20.26 |
| 10.14 | 20.28 | 13.84 |
| 11.65 | 23.31 | 15.90 |
| 9.40 | 18.80 | 12.83 |
| 10.16 | 20.32 | 13.86 |
| 10.34 | 20.67 | 14.10 |
| 10.05 | 20.10 | 13.71 |
| 10.65 | 21.29 | 14.52 |
| 9.45 | 18.90 | 12.89 |
| 6.10 | 12.20 | 8.32 |
| 9.62 | 19.24 | 13.12 |
| 9.64 | 19.28 | 13.15 |
| 5.43 | 10.86 | 7.41 |
| 10.08 | 20.16 | 13.75 |
| 10.58 | 21.16 | 14.44 |
| 9.16 | 18.31 | 12.49 |
| 5.55 | 11.11 | 7.58 |
| 11.51 | 23.03 | 15.71 |
| 10.09 | 20.19 | 13.77 |
| 22.88 | 45.76 | 31.21 |
| 19.46 | 38.93 | 26.55 |
| 17.84 | 35.68 | 24.34 |
| 10.53 | 21.05 | 14.36 |
| 11.40 | 22.79 | 15.55 |
| 5.33 | 10.66 | 7.27 |
| 11.65 | 23.30 | 15.89 |
| 10.71 | 21.41 | 14.61 |
| 5.41 | 10.83 | 7.39 |
| 9.93 | 19.86 | 13.55 |
| 10.16 | 20.31 | 13.86 |
| 6.65 | 13.31 | 9.08 |
| 11.54 | 23.08 | 15.75 |
| 9.61 | 19.21 | 13.10 |
| 11.84 | 23.68 | 16.15 |
| 5.09 | 10.18 | 6.94 |
| 10.45 | 20.91 | 14.26 |
| 10.28 | 20.56 | 14.02 |
| 6.12 | 12.25 | 8.35 |

| | | |
|---|---|---|
| 6.15 | 12.29 | 8.38 |
| 6.23 | 12.47 | 8.50 |
| 6.18 | 12.37 | 8.44 |
| 8.53 | 17.07 | 11.64 |
| 5.58 | 11.16 | 7.61 |
| 6.95 | 13.89 | 9.48 |
| 9.10 | 18.19 | 12.41 |
| 8.42 | 16.84 | 11.49 |
| 20.23 | 40.47 | 27.60 |
| 9.00 | 17.99 | 12.27 |
| 24.52 | 49.04 | 33.45 |
| 8.74 | 17.47 | 11.92 |
| 23.78 | 47.55 | 32.44 |
| 21.50 | 43.00 | 29.33 |
| 11.98 | 23.97 | 16.35 |
| 11.40 | 22.79 | 15.55 |
| 8.62 | 17.24 | 11.76 |
| 9.29 | 18.57 | 12.67 |
| 22.98 | 45.96 | 31.35 |
| 10.82 | 21.64 | 14.76 |
