## Supplementary figures and images for "MorphoHub: A Platform for Petabyte-Scale Multi-Morphometry Generation"

### Supplementary Fig 1

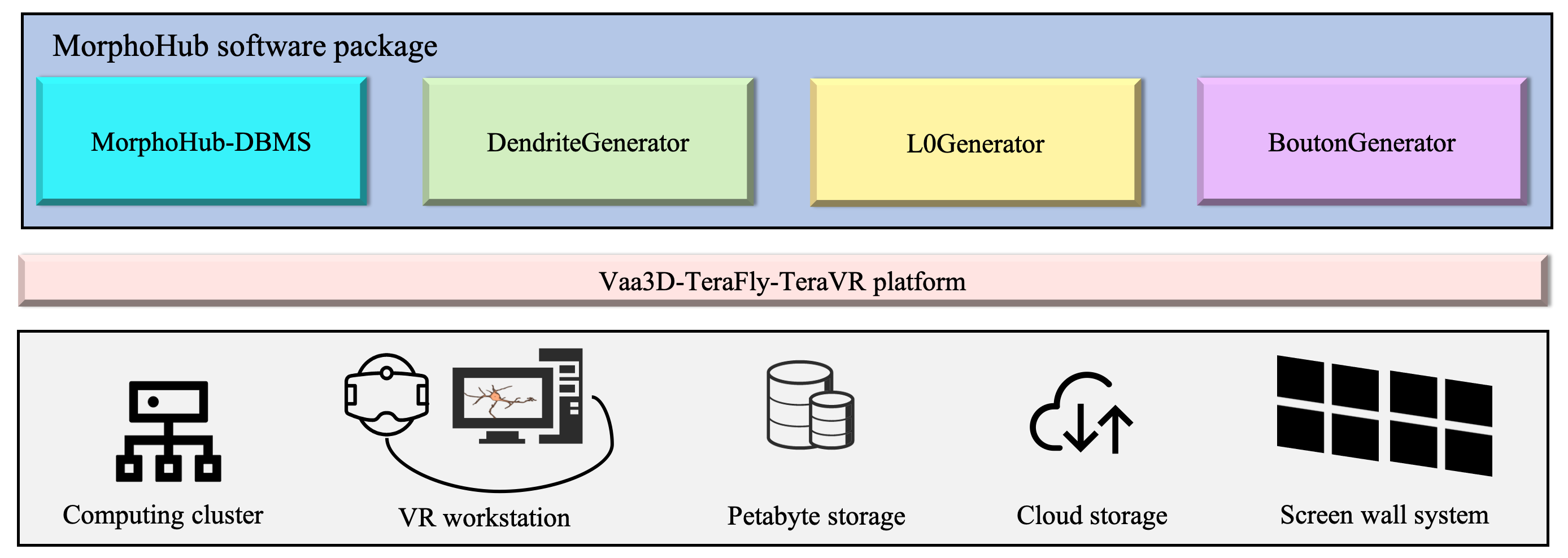

### Supplementary Fig 2

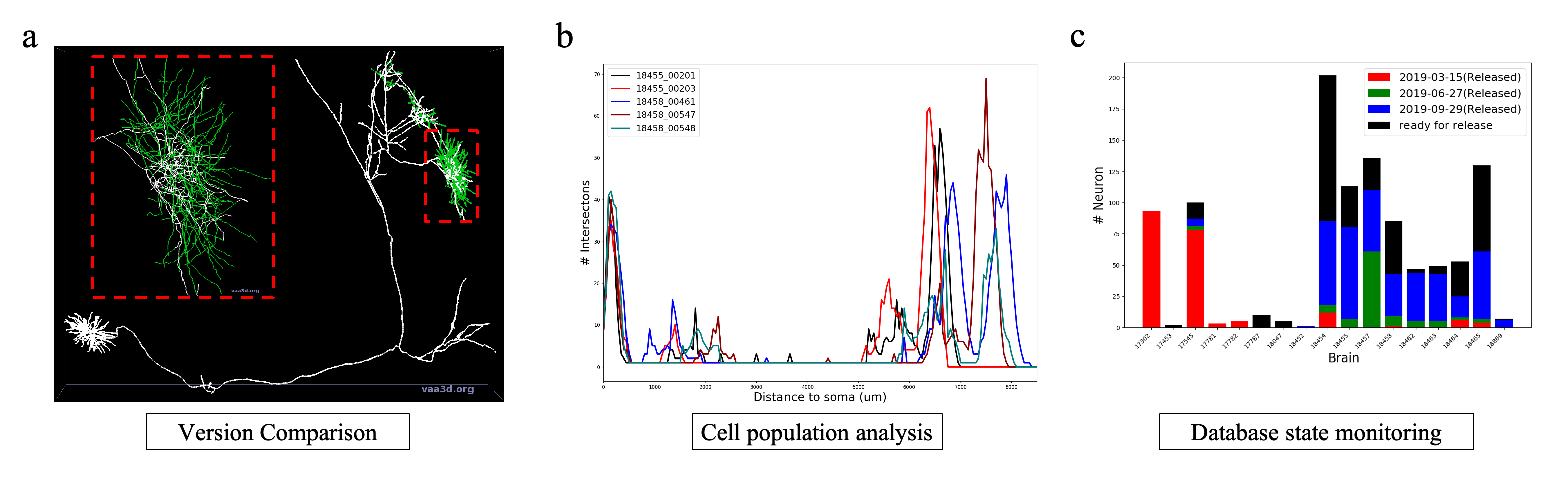

### Supplementary Fig 3

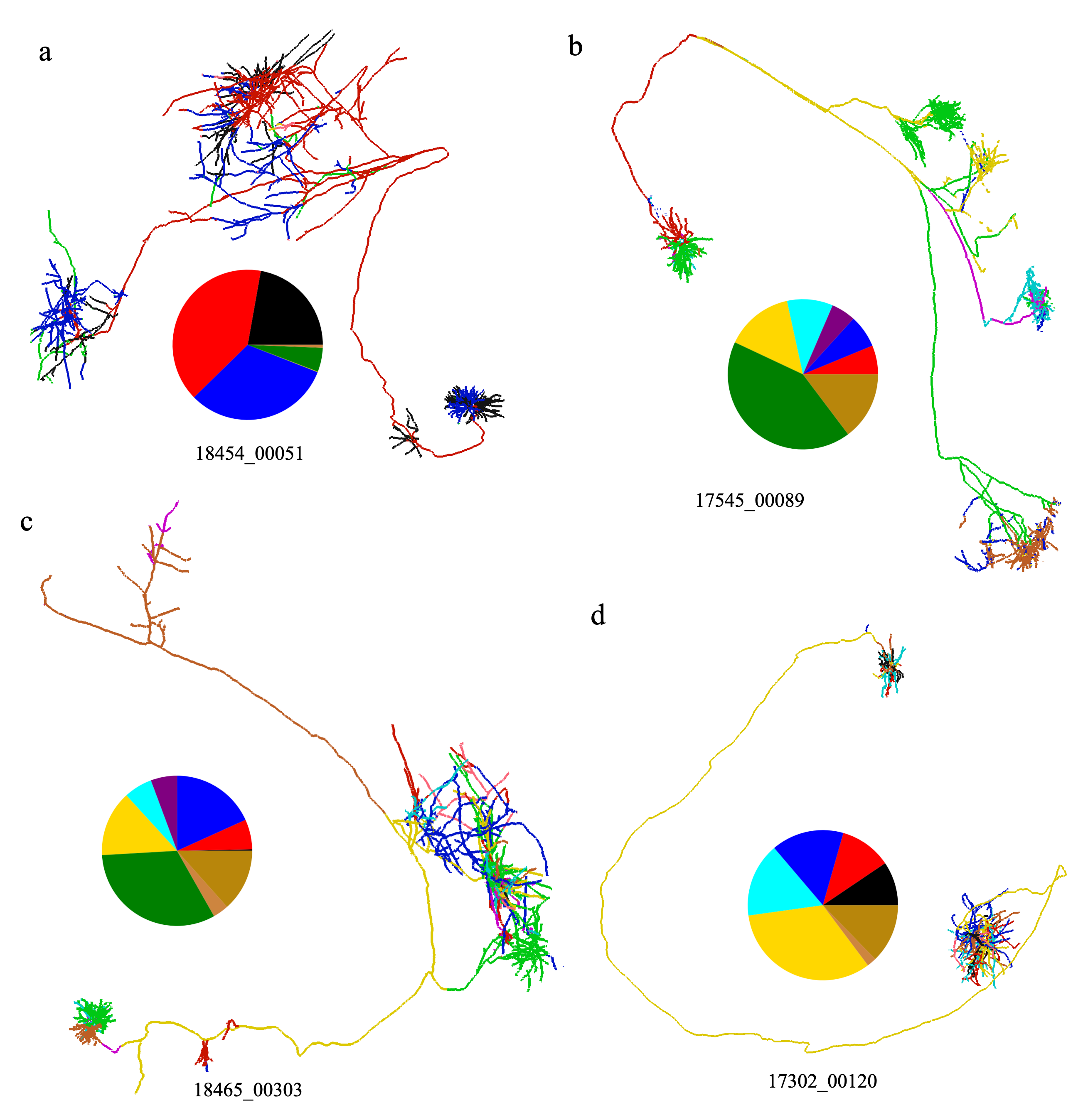

### Supplementary Fig 4

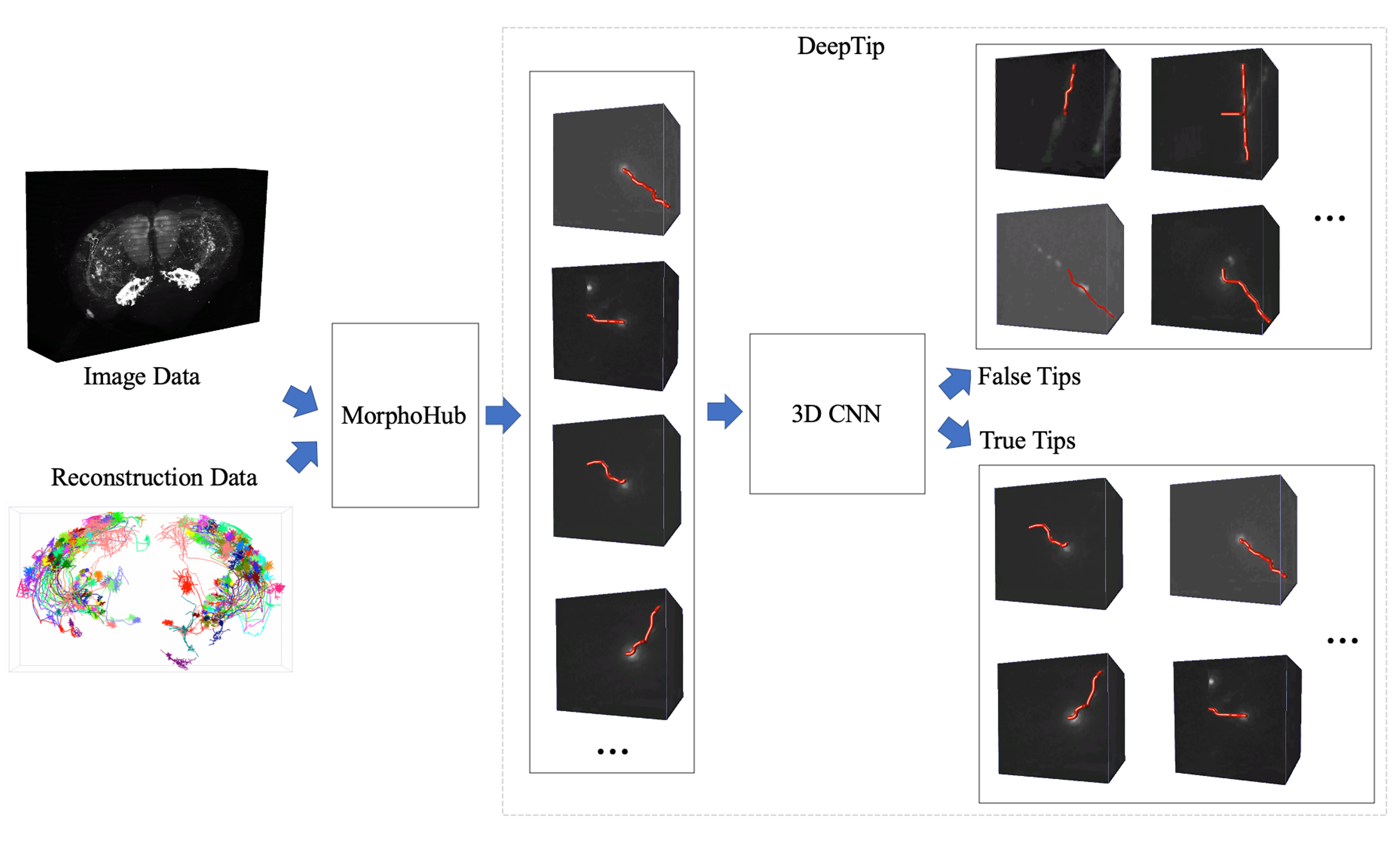
